# Endothelial decorin is increased by ageing and, induces inflammation and diastolic dysfunction in the heart

**DOI:** 10.1101/2025.02.14.638240

**Authors:** Guillermo Luxán, Timm Winkelmeier, Colin Bodemer, Büşra Nur Toğru, Mariana Shumliakivska, Marion Muhly-Reinholz, Ariane Fischer, Mariano Ruz-Jurado, David John, Wesley T. Abplanalp, Stefanie Dimmeler

**Author notes:** Corresponding author, Guillermo Luxán, Goethe University Frankfurt Theodor Stern Kai 7, 60590 Frankfurt; Germany. Equal contribution (Timm Winkelmeier, Colin Bodemer).

## Abstract

**Aims:** Cardiovascular disease is the leading cause of death in the European Union and aging is one of its major risk factors resulting in the progressive deterioration of the cardiac structures and function. Here, we have combined single-nucleus-RNA-sequencing, imaging, and molecular and cell biology approaches to explore the maladaptive signals that drive cardiac ageing.

**Methods and results:** Single-nucleus-RNA-sequencing analysis of young (3 months) and old (18 months) murine hearts revealed that the expression of decorin, a secreted proteoglycan expressed in the extracellular matrix of endothelial cells, is induced by ageing. Decorin treatment via osmotic mini-pump induced diastolic dysfunction and a pro-inflammatory environment in the myocardium characterized by increased infiltration of immune cells, increased expression of IL- 1β in endothelial cells and microvascular leakage in 3 months old mice. In vitro, decorin treatment induces cardiomyocyte hypertrophy, the expression of different pro-inflammatory cytokines like *IL1B* in endothelial cells, and compromises the endothelial barrier function.

**Conclusions:** Together, our results identify decorin as a novel player contributing to cardiac aging and disease. Decorin contributes to the age-related structural and functional dysfunction of the heart by inducing a pro-inflammatory environment in the myocardial microvasculature, a hallmark of cardiac ageing.

**Translational perspective:** Ageing is a major risk factor of cardiovascular disease and the molecular and cellular mechanisms that drive this process have not been completely described. The data presented here identifies decorin as a novel player contributing to systemic inflammation and microvascular dysfunction, two hallmarks of ageing. Although, because of its role regulating TGF-β signalling, decorin has been proposed for anti-fibrotic therapies, the pro-inflammatory effects observed on the cardiac microvasculature should be taken into account for the employment of decorin as an antifibrotic agent to treat disease associated cardiac fibrosis.

## Introduction

Cardiovascular diseases (CVD) are the leading cause of death in the EU accounting for 37% of all deaths ^1,2^. Ageing is a major risk factor in CVD and it leads to the progressive deterioration of cardiac function and structure ^3,4^. Cardiac ageing is characterized by the presence of fibrosis, cardiomyocyte hypertrophy, chronic inflammation and vascular dysfunction ^5^. Furthermore, cardiac ageing is also characterized by the increased prevalence of left ventricular hypertrophy, decline in diastolic function with relatively preserved systolic function at rest but reduced exercise capacity ^3^. Although the phenotypes that arise from cardiac aging have been well described, the molecular mechanisms and maladaptive signals that drive cardiac pathologies are just beginning to be revealed.

The cardiac extracellular matrix is altered during ageing. Not only there is increased collagen deposition and cross-linking, the degradation capacity of the extracellular matrix can also be increased due to the increased expression of different matrix metalloproteases ^6^. Moreover, the cardiac extracellular matrix plays a crucial role controlling heart biology and regeneration ^7,8^. Endothelial secreted extracellular matrix components are considered to be angiocrine agents for their ability to refine the affinity of adhesion molecules or to expose growth factors embedded within the matrix. Angiocrine signals are known to play active roles in cardiac remodelling ^9^. Decorin is a small leucine rich proteoglycan that is secreted into the extracellular matrix ^10^. Decorin is originally known to be involved in collagen fibrillogenesis in the extracellular matrix and obtained its name because it “decorates” the collagen fibrils ^11,12^. Beyond its structural role, decorin has been described to be involved in the regulation of the signal transduction of different pathways involved in cell growth, differentiation, proliferation, adhesion, migration and inflammation ^13^. In the present study, we report that decorin is upregulated specifically in aged endothelial cells in the heart. In order to evaluate the impact of decorin to the heart we have treated young wild type mice with decorin for four weeks and observed that decorin induces cardiomyocyte hypertrophy, inflammation, vascular dysfunction and diastolic dysfunction, hallmarks of cardiac ageing.

## Methods

### Mice

Male and female C57Bl/6J mice were used in this study. Mice were purchased by Janvier (Le Genest Saint-Isle, France). The following transgenic mice were used: decorin fl/fl ^14^ and *Cdh5- CreERT2* ^15^. Endothelial-specific decorin knockout (*decorin^ΔEC^*) was generated by breeding *decorin^fl/fl^* mice with *Cdh5-CreERT2* mice. Decorin deletion was induced in mice aged between 8 and 10 weeks by tamoxifen intraperitoneal injection (20 mg/ml), performed for five consecutive days (100 μl/mouse/day). Homozygous *decorin^fl/fl^* mice were used for animal experiments. We compared animals carrying one copy of *Cdh5-CreERT2* with Cre-negative littermate controls. Experiments were performed in both females and males. Animals with different genotype were kept together in the same cage. Animals were held at 23 °C ambient temperature and 60% humidity in 10 h/14 h light/dark cycle. Animal experiments were performed in accordance with the principles of laboratory animal care as well as German national laws and were approved by the ethics committee of the regional council (Regierungspräsidium Darmstadt, Hessen, Germany) under the animal experiment licenses FU/2007 and FU/2062.

### Cell culture

Before cardiomyocyte isolation hearts from wild type young animals were treated using basal buffer (130 mM NaCl, 5 mM KCl, 0.5 mM NaH2PO4, 10 mM HEPES, 10 mM Taurine, 10 mM Glucose, 10 mM BDM). First the hearts were perfused with basal buffer and 5 mM EDTA and then perfused with basal buffer containing 1 mM MgSO4. The heart was digested with basal buffer plus MgSO4 and 120 U/µl of Collagenase Type II. Digestion was stopped using basal buffer containing 10 % FCS. Cardiomyocyte fraction was enriched by centrifugation 2 minutes at 20 g for three times. Cardiomyocytes are present in the pellet. Cells were cultured in laminin-coated plates using M199 medium containing 25 mM HEPES, 5 mM Creatinine, 2 mM L-Carnitine, 5 mM Taurine, 1% Penicillin / Streptavidin and 10% FCS.

Human cardiomyocytes ventricle type (HCM VT, 36044-15VT, Celprogen) were cultured in collagen-coated flasks containing cell specific serum free media (M3604415, Celprogen). Cells were cultured at 37°C and 5% CO2. Upon reaching the confluency 80-90 %, cells were split using TrypLE™ (12563011, ThermoFischer Scientific). Experiments were performed using cells between passages 2 and 6.

Human cardiac microvasculature endothelial cells (HMVEC-Cs, CC-7030, Lonza) were cultured in EGM-2 MV Microvascular Endothelial Cell Growth Medium-2 Bullet Kit (CC-3202, Lonza). To detach the cells for experiments, reseeding, and passage TrypLE™ (12563011, ThermoFischer Scientific) was used. Experiments were performed using cells at passage 2 and 3. Medium was changed every second day and cells were cultured at 37°C, 5% CO2.

### Decorin treatment

Cells were cultured in cell specific media and 24 h after seeding the medium was changed with medium containing 10 µg/ml recombinant human decorin (143-DE, BioTechne) or recombinant mouse decorin (1060-DE, BioTechne). Decorin treated cells were incubated in cell culture dishes for 72 h and analysed afterwards. For freshly isolated cardiomyocytes the medium containing decorin was refreshed every day.

### RNA-isolation and RT- qPCR

Total RNA was purified from cells using the miRNeasy mini kit (217004; Qiagen), combined with on-column Dnase digestion (Rnase-free Dnase-set; 79254; Qiagen), as described by the manufacturer’s instructions. 500 ng of RNA from each sample were used to synthetize the complementary DNA, using the M-MLV reverse transcriptase (28025013, ThermoFisher Scientific). Real-time qPCR was performed using the Fast SYBR Green master mix (4385617; ThermoFisher Scientific) and the Viia7 – Realtime qPCR System (Life-Technologies). Expression levels were normalised to *GAPDH* and analysed using the 2^-ΔCt^ Method. Primer sequences: *IL1B*: forward, TGAGCTCGCCAGTGAAATGA and reverse, CATGGCCACAACAACTGACG. *IL12:* forward, AAGGAGGCGAGGTTCTAAGC and reverse, GAGCCTCTGCTGCTTTTGAC. *GAPDH*: forward, TCGAGTCAGCCGCATCTTCTTT and reverse, ACCAAATCCGTTGACTCCGACCTT.

### Immunocytochemistry

Cells were fixed with 4% PFA for 10 min. After washing twice for 5 min with PBS, cells were permeabilized with 0.1% Triton X-100 in PBS for 15 min. Cells were blocked with 5% donkey serum in PBS for 60 min at RT. Cells were stained with Phalloidin Alexa Fluor 488 (A12379, Invitrogen) (1:100) in the same blocking solution overnight at 4°C. Nuclei were counterstained using DAPI (D9542; Sigma) (1:1000). After washing twice with PBS, cells were mounted with Fluoromount-G (Invitrogen). Stainings were imaged in a Leica SP8 and Stellaris confocal inverted microscopes. Quantification was done using Volocity software version 6.5 (Quorum Technologies).

### Cytokine array

HMVEC-Cs cells were cultured and treated with decorin as stated above. Afterwards, the cells were washed with PBS and 150 µl Pierce RIPA buffer (89900) (1:100 phosphatase, 1:10 protease) were added to the cells. The cells were rocked at 2-8°C for 30 min and centrifuged at 14,000 x g for 5 min at 4°C. Afterwards the supernatant was transferred into a clean tube and the sample protein was quantitated with the DC Protein-Assay Kit II (Bio-Rad, #5000112). The cytokine array was performed according to manufactureŕs instruction using the Proteome Profiler Human Cytokine Array Kit (ARY005B; R&D Systems). The quantification was performed using Image J Fiji Protein Array Analyzer.

### Western Blot

Hearts from 3- and 18-month-old mice were homogenized and lysed in RIPA buffer (R0278, Sigma-Aldrich) supplemented with protease (1:50, P8340, Sigma-Aldrich) and phosphatase inhibitors (1:100, P5726, Sigma-Aldrich). Protein concentration was measured using the DC Protein Assay Kit II (5000112, Bio-Rad) following manufacturer’s instructions. After denaturation at 95°C, proteins were loaded on pre-casted gradient gels (4561094/5678094, Bio-Rad) and separated by SDS-PAGE in 1X Tris/Glycine/SDS Buffer (1610732, Bio-Rad). Transfer onto nitrocellulose membrane (10600006, Cytiva) was performed in 1X Tris/Glycine Buffer (1610734, Bio-Rad) using the Trans-Blot® Semi-Dry system (1703940, Bio-Rad). Membranes were blocked in 5% milk (sc-2324, Santa Cruz) and probed with primary antibodies overnight at 4°C followed by incubation with ECL-HRP-linked secondary antibody (1:1000, NA934V, Amersham). Primary antibodies used: rabbit anti-decorin (1:1000, ABT274, Merck), rabbit anti-GAPDH (1:1000, 2118S, Cell Signalling). Proteins were detected by chemiluminescence using HRP Substrate (WBKLS0500, Millipore) in a ChemiDoc Touch imaging system (Bio-Rad). Bands were quantified by densitometry with ImageJ (version 1.52p).

### Decorin treatment with Osmotic mini pumps

Recombinant mouse decorin (1060-DE, BioTechne) was diluted in PBS at a concentration of 100 µg/ml ^16,17^ and delivered using Osmotic mini pumps (1004, Alzet) at a rate of 25 µl/h ^18^.

### Echocardiography

Echocardiography was performed using a Vevo3100 (FUJIFILM VisualSonics, Inc, Toronto, Canada).

### Immunohistochemistry

Immunohistochemistry was performed as previously ^19,20^. Hearts were dissected in cold PBS and fixed overnight in 4% PFA at 4°C. Samples were then cryo-protected with a series of consecutive overnight washes in PBS containing increasing concentration of sucrose (Sigma, S0389) (10%, 20% and 30%) and finally embedded in embedding solution (15% sucrose, 8% gelatin (Sigma, G1890) and 1% polyvinylpyrrolidone (pvp; Sigma, P5288) in PBS). When embedding solution solidified, tissue blocks were frozen at −80°C. 50 µm thick cryosections were produced using a Leica CM3050S cryostat. Cryosections were allowed to dry overnight at room temperature and then stored at −20°C until the staining procedure. For the WGA staining, the tissue was embedded in paraffin. In this case, before the starting of the immunohistochemistry procedure the samples were deparaffined and rehydrated. In case of the other stainings, before starting the immunohistochemistry procedure, samples were allowed to acquire room temperature. Slides containing the samples were then rehydrated in PBS by washing them two times for 5 minutes and then permeabilized with PBS containing 0.3% Triton X-100 by washing them three times for 10 minutes. Tissue was blocked for one hour in freshly prepared blocking solution (3% BSA, 0.1% Triton X-100, 20 mM MgCl2, and 5% donkey serum in PBS). Primary antibodies were incubated overnight at 4°C in the blocking solution. On the following day, the primary antibodies were washed four times 5 min in PBS. Secondary antibodies, together with DAPI, were incubated for one hour at room temperature in PBS containing 5% BSA. Immunostainings were imaged in a Leica SP8 and Stellaris confocal inverted microscope.

Primary antibodies: Rabbit anti-NG2 (AB5320; Millipore) (1:100), WGA-Alexa Fluor 647 (2538557; Invitrogen) (1:100), biotinylated Isolectin-B4 (B-1205.5; Vector Laboratories) (1:25), rabbit anti-decorin (AbT274, Merck) (1:50), rabbit anti-mouse serum Albumin (Ab19196, Abcam) (1:100), rat anti-Ter119 (MAB1125, R&D Systems) (1:20), mouse anti-IL1β (12242S, Cell signaling) (1:100), rat anti-CD45 (Ab25386, Abcam) (1:50), rabbit anti-CD68 (97778S, Cell signaling) (1:100). Secondary antibodies: Donkey anti rabbit Alexa Fluor 488 (A21206, Invitrogen) (1:200), donkey anti rabbit Alexa Fluor 555 (A32794, Invitrogen) (1:200), donkey anti rabbit Alexa Fluor 647 (A32795, Invitrogen) (1:200), donkey anti rat Alexa Fluor 594 (A48271, Invitrogen) (1:200), donkey anti mouse Alexa Fluor 647 (A32787, Invitrogen) (1:200), donkey anti goat Alexa Fluor 647 (A32849, Invitrogen) (1:200), Streptavidin Alexa Fluor 488 (S11223; Invitrogen) (1:200) and Streptavidin Alexa Fluor 647 (S32357; Invitrogen) (1:200). Nuclei were counterstained using DAPI (D9542; Sigma) (1:1000).

Stainings were quantified using Volocity Software 6.5.1 (Quorum Technologies).

### Sirius red staining

To analyse collagen deposition in the heart, we performed Sirius Red staining on paraffin sections. Fixed hearts (as described above) were embedded in paraffin. After deparaffinization and rehydration, the slides were stained with 0.1% Picro Sirius Red solution prepared using Sirius Red F3BA (1A280, Waldeck GmbH) in an aqueous solution of Picric Acid (6744, Sigma Aldrich) following standard protocol. Slides were mounted with Pertex mounting medium (00801-EX, HistoLab) and images were acquired using a Nikon Eclipse Ci microscope with a 40X objective. Positive area was measured using ImageJ (NIH, version 1.53a) and normalised to the total tissue area.

### Single-nucleus-RNA-sequencing

The single-nucleus-RNA-sequencing data set generated in the paper by Vidal, Wagner et al. ^21^, was used in this study for the young and old mice comparison. For this study, three 3-month-old males and three 18-months-old males were used.

### Gene ontology

Gene ontology analysis was performed using Gene Ontology Database with the R package clusterProfiler (4.2.2).

### Permeability assay

HMVEC-C (CC-7030, Lonza) cells were seeded on a 6-well plate. After 4h medium was changed to medium containing 10 µg/ml recombinant human Decorin (143-DE, Biotechne) or the same volume of PBS and incubated for 3 days. After 3 days, supernatants were collected. Meanwhile fresh naive HMVEC-C cells were seeded on top of a fibronectin (1:200) coated cell culture insert (ThinCert, 1 µm pore diameter, 24-well, Greiner Bio-One) and the collected supernatants were added on top of the cell culture inserts and incubated for 24 h. 60 ng/mL VEGF (SRP4369, Sigma) treatment 1 hour before the end of the experiment was used as positive control. Before measurement, the 24-well and inserts were washed twice with PBS. Then, 250 µL Opti-MEM containing 10 µg/ml 70 kDa FITC-Dextran (46945, Sigma) was added to the chamber and 850 µL Opti-MEM to the bottom well and incubated for 1 h at 37°C. Fluorescence (λex = 493 nm, λem = 518 nm) in the bottom well was measured with a GloMax-Multi+Detection System (Promega).

### Statistics

GraphPad’s 9 (Prism) software was used for statistical analysis of all experiments but single-nucleus-RNA-seq. Shapiro–Wilk normality test was used to test data before comparison. Unpaired, Student’s t-test was used for comparison between two groups according to their distribution. For comparisons between more groups, one-way ANOVA followed by Tukey’s multiple comparisons test was employed. Data is presented in scatter plots with mean ± standard error of the mean (SEM) unless indicated otherwise.

## Results

### 1. Decorin expression in endothelial cells is increased by ageing

Gene ontology analysis of the differentially expressed genes obtained from single nuclei RNA sequencing data comparing 3-month-old (3M) and 18-month-old (18M) mice ^21^ across all cell types revealed that the cellular compartment to which most of these genes belong is the extracellular matrix and the collagen-containing extracellular matrix. Furthermore, other external components of the cell like the external encapsulating structure or the basement membrane are also upregulated (**Figure 1A**). Detailed study of the genes comprised in these categories showed that *Dcn* was upregulated specifically in aged endothelial cells (**Figure 1B**). *Dcn* is also expressed in cardiac fibroblasts (**Supplementary Figure 1A**), but its expression is not regulated in the aged heart (**Supplementary Figure 1B**). Furthermore, endothelial cells are the most predominant cell type in the heart and thus, gene expression alterations in this cell type may play a bigger role in physiologic function and response to injury than in other cell types ^22^. Transcriptome analysis of endothelial cells isolated from aged hearts ^23^ further confirmed this upregulation (**Figure 1C**). Histological analysis of young and old hearts showed an accumulation of decorin in the aged heart (**Figure 1D, E**) and finally, western blot analysis of total heart lysates further confirmed the increase of decorin expression in the heart (**Figure 1F, G**). Taken together, these observations allow us to hypothesize that the increased accumulation of decorin in the aged myocardium is due to the increased gene expression in endothelial cells.

**Figure 1.**
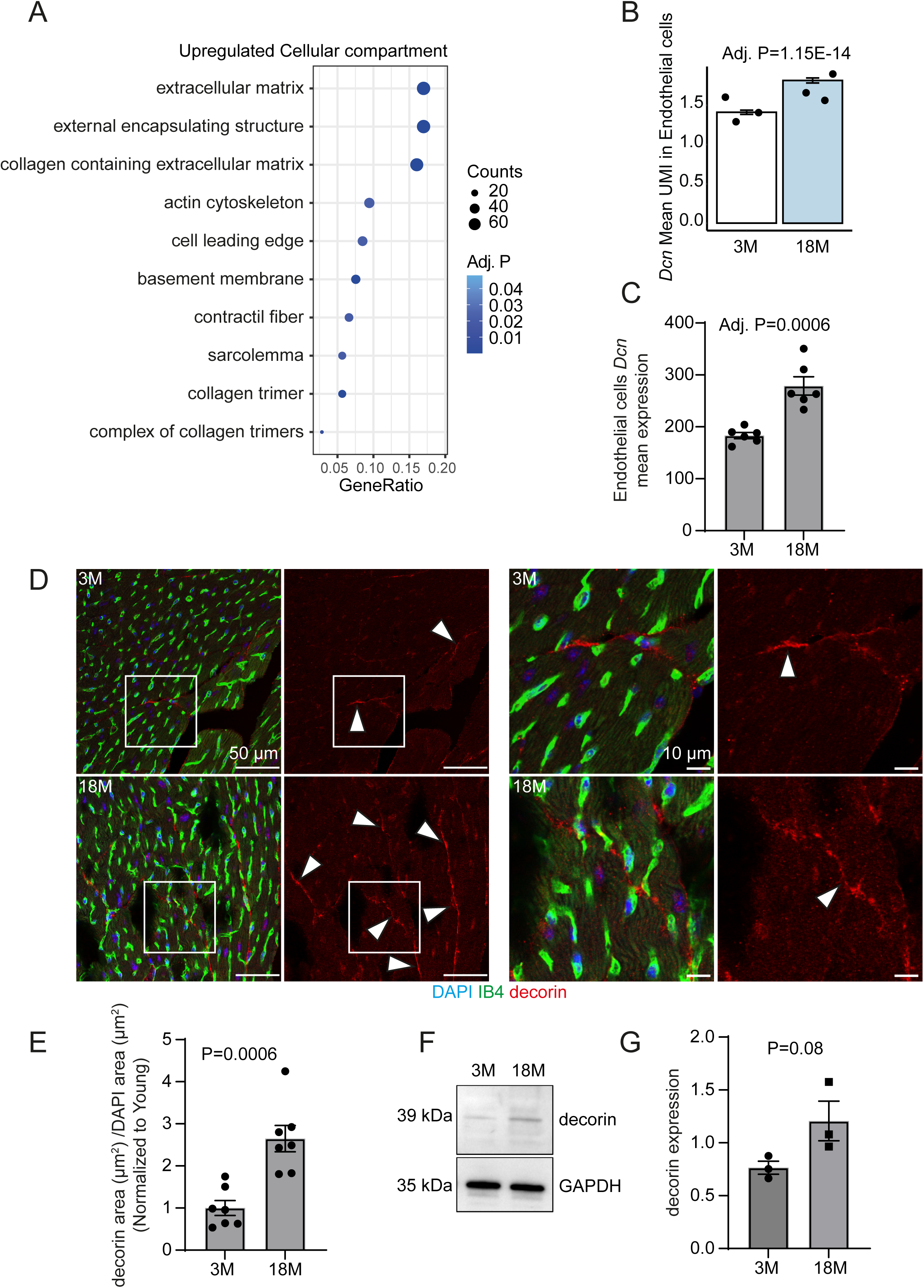
Decorin is upregulated in the old heart. (**A, B**) Single nucleus RNA sequencing. (A) Gene ontology analysis representation of the ten most enriched cellular compartments in the aged heart. (**B**) Bar chart showing increased *Dcn* expression in old and young endothelial cells (N=3). P-value calculated by the Wilcoxon-Rank Sum test. (**C**) Bulk RNA sequencing. Bar graph showing increased *Dcn* mean expression in isolated endothelial cells from young and old hearts (N=6). Data are shown as mean ± SEM. P-value was calculated by unpaired two-tailed Student’s t-test. (**D**, **E**) Immunostaining of cross sections through the left ventricle showing increased expression of decorin in the old hearts (N=7). Data are shown as mean ± SEM. P-value was calculated by unpaired two-tailed Student’s t-test. (**F**, **G**) Western blot of protein lysates isolated from 3M and 18M hearts. Data are shown as mean ± SEM. P-value was calculated by unpaired two-tailed Student’s t-test.

### 2. Decorin induces diastolic dysfunction in the heart

To study whether decorin could induce age related features in the heart, we implanted osmotic mini pumps containing decorin or PBS on wild type 3-month-old animals and followed them for four weeks (**Figure 2A**). Similar to what we observed in the aged mice, decorin accumulated in the interstitial space between capillaries and cardiomyocytes in the heart of treated mice four weeks after starting the treatment (**Figure 2B, C**) indicating the success of decorin delivery. Echocardiography analysis (**Figure 2D** and **Supplementary Figure 2**) of the hearts after decorin treatment showed no effects on the thickness of the ventricular septum, and no effect on the systolic function of the heart measured by ejection fraction (**Figure 2D**). Nevertheless, the treatment with decorin induced an increase of the mitral E/E’ ratio, a sign of diastolic dysfunction (**Figure 2D**). Reduced diastolic function but preserved systolic function are characteristics of the aged heart ^24^. Further characterization of the phenotype induced by decorin treatment revealed no effects on cardiomyocyte hypertrophy *in vivo* (**Figure 2E, F**). Nevertheless, treating both, freshly isolated murine cardiomyocytes and commercially available human primary cardiomyocytes of ventricular origin, with decorin for three days was sufficient to induce cardiomyocyte hypertrophy *in vitro* (**Supplementary Figure 3A, B**). Sirius red staining of the myocardium revealed that decorin treatment did not induce fibrosis in the treated animals (**Figure 2G, H**). Although fibrosis is a hallmark of ageing ^25^, decorin has been already shown to possess antifibrotic capabilities ^26,27^.

**Figure 2.**
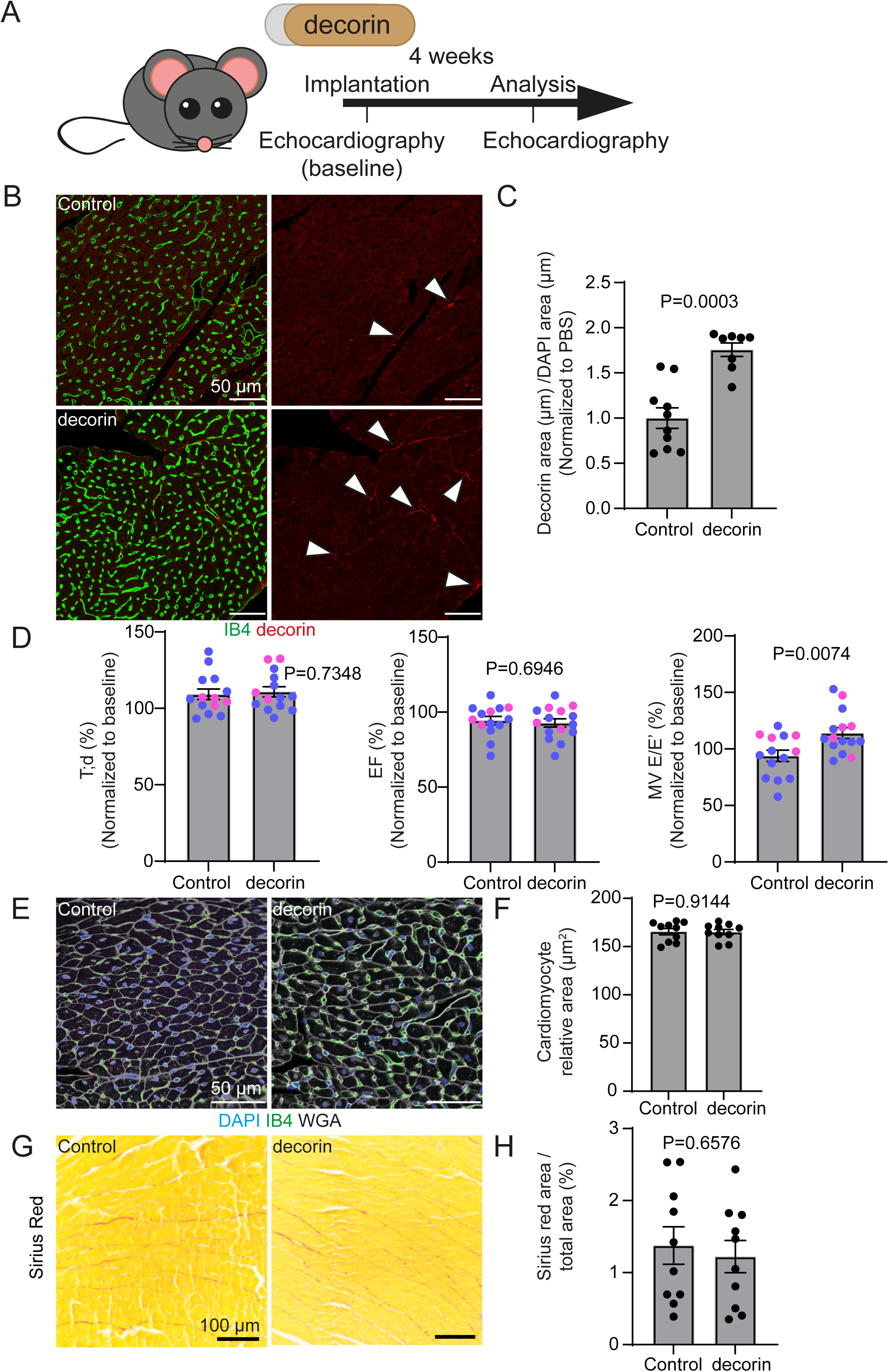
Decorin treatment induced diastolic dysfunction *in vivo*. (**A**) Scheme of the experimental design. (**B**, **C**) Immunostaining of cross sections through the left ventricle showing increased presence of decorin in the treated hearts (N=10). Data are shown as mean ± SEM. P- value was calculated by unpaired two-tailed Student’s t-test. (**D**) Echocardiographic analysis of animals after decorin treatment. (N=14 for Control and N=15 for decorin). Colour indicates the sex of the animal, blue represents male, pink represents female. Data are shown as mean ± SEM. P-value was calculated by unpaired two-tailed Student’s t-test. (**E**, **F**) Immunostaining of cross sections through the left ventricle. Decorin treatment does not affect cardiomyocyte size (N=10). Data are shown as mean ± SEM. P-value was calculated by unpaired two-tailed Student’s t-test. (**G**, **H**) Sirius red staining of heart cross sections. Decorin does not increase collagen deposition in the heart (N=10). Data are shown as mean ± SEM. P-value was calculated by unpaired two- tailed Student’s t-test.

Ageing and heart failure are well known to be associated with microvasculature impairment ^28,29^. Detailed analysis of the microvasculature in the treated hearts revealed that decorin induced increase of the capillary perimeter without altering the vascular density in the heart (**Figure 3A- C**). Cardiac capillary perimeter enlargement is feature of the aged murine heart ^30^. Furthermore, the vascular integrity was compromised upon treating the animals with decorin as the levels of plasma Albumin were increased in the myocardium (**Figure 3D, E**). Moreover, extravasated Albumin could be observed in the interstitial space of the myocardium (**Figure 3D, F, arrowheads**). The increased microvascular leakage was accompanied by increased immune cell infiltration. Both the general immune cell marker CD45 and the macrophage marker CD68 are increased in the myocardium of the decorin treated hearts (**Figure 3G-I**). Despite these effects on the cardiac microvasculature, we could not observe any effect on the pericyte coverage of the capillaries (**Supplementary Figure 4**).

**Figure 3.**
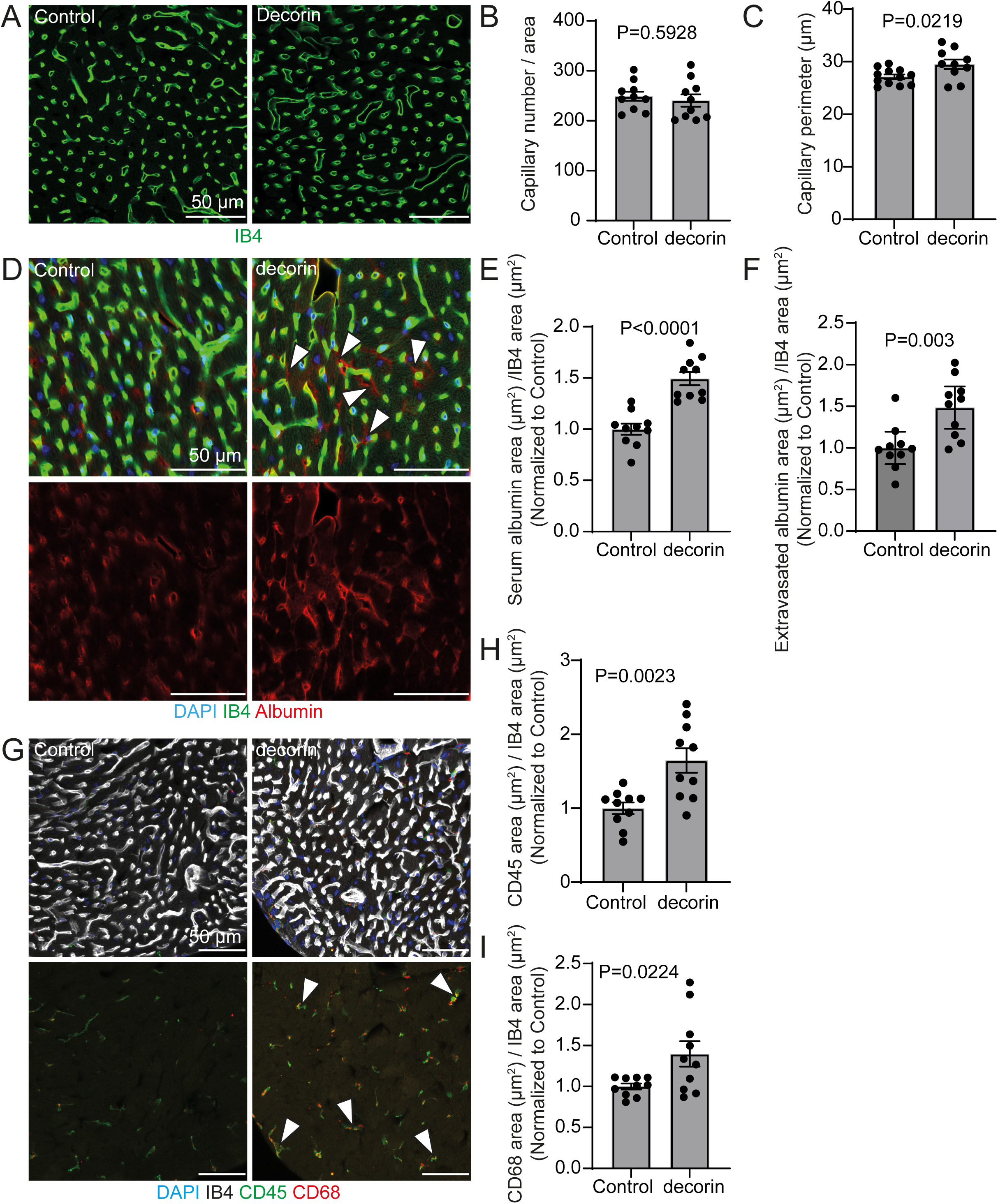
Decorin treatment induced vascular dysfunction. (**A**-**I**) Immunostaining of heart cross sections (N=10). (**A**-**C**) Decorin induces capillary dilation in the heart. (**D**, **F**) Decorin treatment induces vascular leakage as detected by extravasated Albumin and (**G-I**) immune cell infiltration. Data are shown as mean ± SEM. P-value was calculated by unpaired two-tailed Student’s t-test.

### 3. Decorin treatment induces a pro-inflammatory environment in the heart

Chronic low-grade inflammation is characteristic of aged individuals and can cause vascular dysfunction ^31^. The increased number of immune cells in the myocardium suggested that in our model, decorin induced a pro-inflammatory environment in the myocardium *in vivo*. To test this, we analysed the expression of the pro-inflammatory cytokine IL-1β in capillary endothelial cells.

Decorin treatment did indeed induce the expression of IL-1β in capillary endothelial cells (**Figure 4A, B**). We then tested whether decorin, independent of other age-related cues would induce vascular inflammation *in vitro*. To do so, we treated human cardiac microvasculature endothelial cells (HMVEC-Cs) for three days with decorin (**Figure 4C**). This assay confirmed the increased expression of *IL1B* upon decorin treatment (**Figure 4D**). Furthermore, other pro-inflammatory cytokines like *IL12,* CCL2 and TNFα were also upregulated upon decorin treatment (**Figure 4E**- **H and Supplementary Figure 5**). To test whether decorin could directly induce vascular dysfunction similar to the one we had observed *in vivo*, we tested HMVEC-C cell permeability. To test our hypothesis that decorin induced inflammation would impair barrier function, we treated HMVEC-C cells with decorin for three days and collected the conditioned supernatant. Then, we treated naïve HMVEC-C cells seeded on a boyden chamber either with the supernatant obtained from decorin or PBS treated endothelial cells. Indeed, we could detect increased FITC-Dextran permeability when the cells were treated with the supernatant from endothelial cells that have been previously treated with decorin. The permeability increase was similar to the one observed when the cells were treated with the positive control VEGF ^32^, confirming the role of decorin inducing endothelial barrier dysfunction (**Figure 4I**). Decorin can act as a damage-associated molecular patterns (DAMP) molecule serving as a ligand of TLR2/4 and triggering inflammatory responses ^33,34^. TLR2 activation has already been shown to induce the production of IL-1β ^35,36^ This leads us to hypothesize that in endothelial cells, decorin may activate TLR2, which in turn drives the expression of *IL1B* and subsequently causes vascular leakage To test this hypothesis, we depleted TLR2 from endothelial cells using siRNA (**Figure 4J, K**). Indeed, this treatment reduced the expression of IL1B upon decorin treatment (**Figure 4L**). Moreover, inflammation and thrombosis are two well connected entities ^37^. To test whether decorin induced inflammation can cause thrombosis in the myocardium capillaries we perform immunostainings against Ter119, a marker of erythrocytes, in the myocardium of the decorin treated animals. Indeed, we could detect an increased number of erythrocytes trapped in the capillaries of the treated hearts (**Supplementary Figure 6**). Finally, to test whether endogenous endothelial decorin play a role in inflammation and vascular leakage, we bred mice bearing a conditional *Dcn* loss-of-function allele ^14^ with *Cdh5-CreERT2* transgenic animals, which express tamoxifen-inducible Cre recombinase specially in endothelial cells ^15^ (**Figure 5A**). The deletion of endogenous endothelial decorin (**Figure 5B**) reduced the expression of IL-1β by capillary endothelial cells (**Figure 5C, D**). We also analysed whether the deletion of decorin in endothelial cells would have an effect on the accumulation of serum albumin in the myocardium, but we did not observe any difference between the transgenic and control animals (**Figure 5E, F**). Nevertheless, the observed reduction of IL-1β supports our previous observations suggesting that indeed, decorin contributes to an inflammatory phenotype in the capillary endothelial cells.

**Figure 4.**
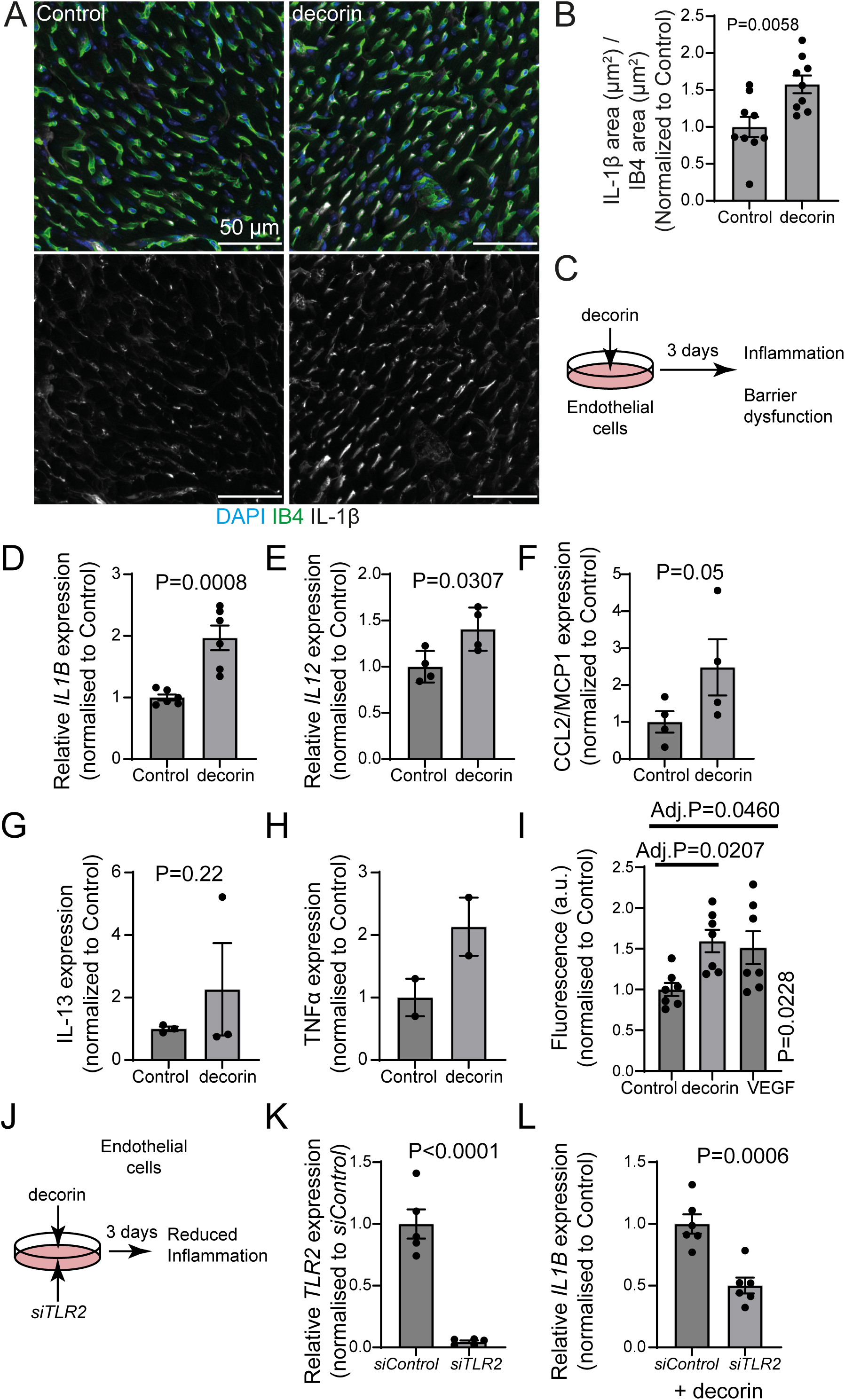
Decorin induced inflammation in the heart. (**A**, **B**) Immunostaining of heart cross sections (N=10). Decorin induces IL-1β in the capillaries in the heart. Data are shown as mean ± SEM. P-value was calculated by unpaired two-tailed Student’s t-test. (**C**) Scheme of the experimental decorin treatment. (**D**, **E**) qPCR analysis of gene expression in HMVEC-Cs upon decorin treatment. The expression of the pro-inflammatory genes *IL1B* (N=6) (**D**) and *IL12* (**E**) (N=4) are upregulated in treated cells. Data represented as mean ± SEM. P-value was calculated by unpaired two-tailed Student’s t-test. (**F**-**H**) Cytokine array expression analysis HMVEC-Cs upon decorin treatment. Data represented as mean ± SEM. P-value was calculated by unpaired one-tailed Student’s t-test. (**F**) Expression of CCL2/MCP1. (N=4). (**G**) Expression of IL-13. (N=3). (**H**) Expression of TNFα. (N=2). (**I**) Permeability assay of HMVEC-C monolayer upon treatment with the supernatant from decorin treated HMVEC-C. Decorin treatment increased the permeability of the endothelial monolayer to a similar level as VEGF (N=7). P-value was calculated by one way ANOVA followed by Tukey’s multiple comparisons test. (**J**) Scheme of the experimental decorin treatment with *TLR2* knockdown. (**K**, **L**) qPCR analysis of gene expression. Data represented as mean ± SEM. P-value was calculated by unpaired two-tailed Student’s t-test (**K**) TLR2 expression upon siTLR2 transfection. (N=5). (**L**) IL1B expression after TLR2 depletion and decorin treatment. (N=6).

**Figure 5.**
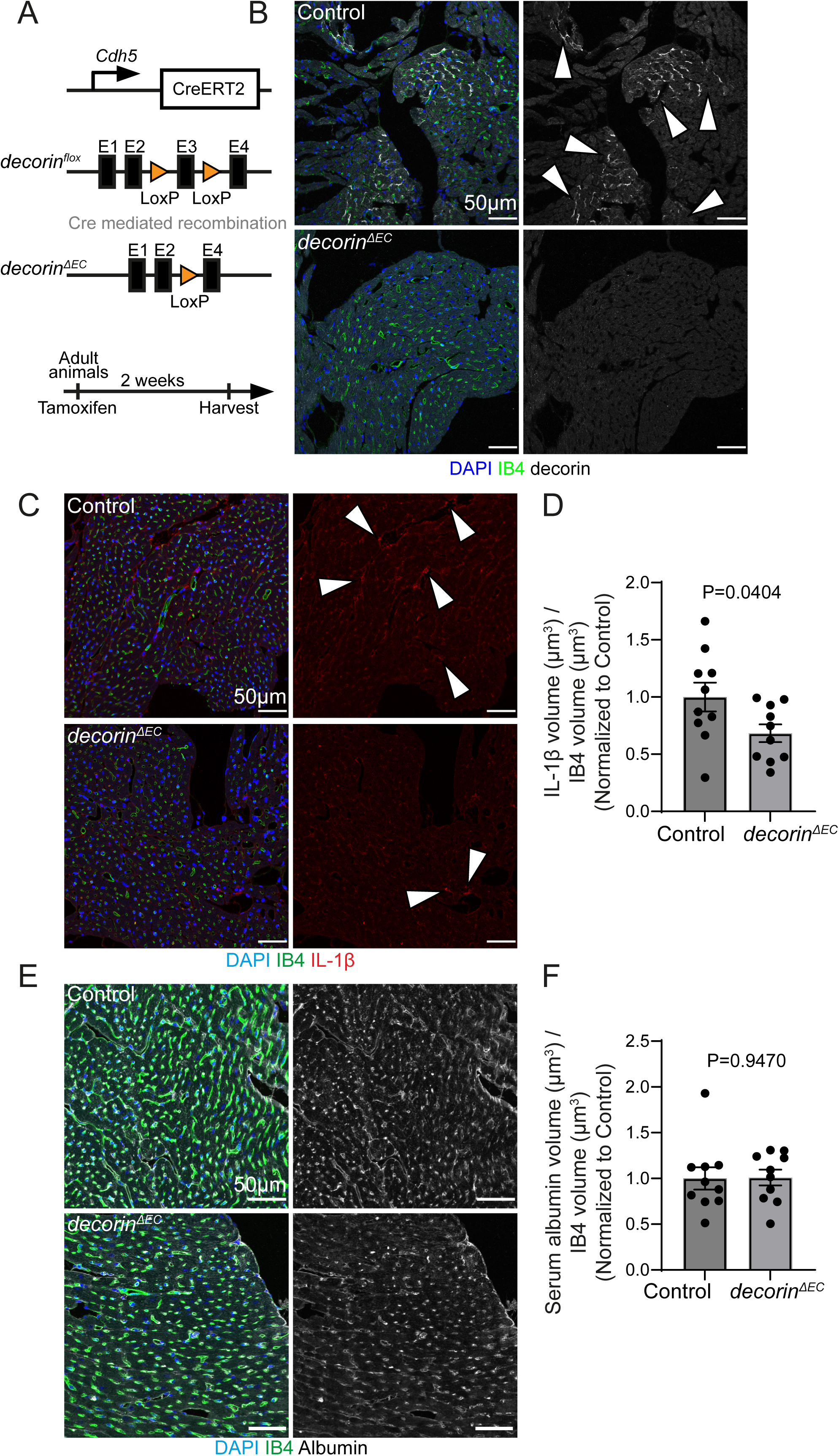
Decorin endothelial deletion rescues IL-1β in the capillaries. (**A**) Scheme of the experimental design. (**B**-**F**) Immunostaining of heart cross sections (N=10). (**B**) Decorin expression. (**C**, **D**) IL-1β expression and (**E**, **F**) serum albumin expression. Data represented as mean ± SEM. P-value was calculated by unpaired two-tailed Student’s t-test.

## Discussion

Here, we show for the first time that the expression of decorin is a sign of cardiac ageing that contributes to cardiac dysfunction. Our *in vitro* and *in vivo* studies have shown that decorin induces different features related to cardiac ageing like cardiomyocyte hypertrophy, vascular inflammation and diastolic dysfunction. The role of the ECM providing the environment and structural support for many cells in the body influences many processes in the body and thus, age related alterations of the ECM might have far reaching consequences for the maintenance of cardiac homeostasis. Decorin expression is linked to ageing in tendons and the genetic deletion of decorin decreased the effects of ageing of tendons ^38^. Decorin was shown to alter collagen fibril structure and alignment in tendons. Decorin induces a reduction in the average fibril diameter ^39^ and in the skin, the absence of decorin leads to a loosely packed collagen network and skin fragility ^40^. We hypothesize that the increased decorin expression in the heart might lead to alterations in the microenvironment where capillaries, cardiomyocytes, and fibroblasts reside. Moreover, similar to our observation, it was already reported that decorin induces cardiomyocyte hypertrophy by regulating the CaMKII-MEF-2 signalling cascade ^41^.

We have furthermore observed that decorin induces increased inflammation in the cardiac capillaries. Consistent with previous findings ^33–36^, our results suggest that decorin acts as a DAMP in endothelial cells and triggers inflammatory responses in a TLR2-dependent manner. Chronic inflammation causes endothelial dysfunction ^42^, and in particular it was shown that IL-1β induces vascular permeability ^43^, and microvascular dysfunction was found to be a major determinant for the development of different cardiac diseases like HFpEF ^29,44,45^. Furthermore, our cytokine array revealed that decorin induces the expression of CCL-2 and TNFα. TNFα is associated to cardiac inflammation, dysfunction and finally heart failure while CCL-2, a small cytokine of the CC chemokine family, is responsible for the recruitment of monocytes to the inflammation and its increased expression may explain the increase of macrophages that we observed in the decorin treated animals. Interestingly, a mouse model of heart failure with preserved ejection fraction (HFpEF) showed similar vascular features to the ones induced by decorin ^46^.

Decorin inhibits fibrosis by blocking TGF-β ^47^ and it has been previously proposed as a potential anti fibrotic therapy ^48,49^. In particular, it has been shown that *Dcn* gene transfer neutralizes TGF-β and attenuates cardiac remodelling after myocardial infarction ^50,51^. Furthermore, local decorin gene therapy reduced the cardiac fibrosis induced by hypertension ^52^. In rats, decorin has been shown to reduce the inflammation related to fibrosis ^53,54^. Nevertheless, we have shown here that endothelial decorin depletion reduces IL-1β expression in the myocardium while decorin treatment induces endothelial inflammation and vascular dysfunction via *TLR2*. We believe that this vascular dysfunction may lead to cardiac dysfunction. These effects on the microvasculature should be considered for the future development of any decorin based therapeutic strategy for cardiac fibrosis as microvascular dysfunction is one of the hallmarks of ageing and different diseases like HFpEF.

In conclusion, our results identified the increase of endothelial decorin as an indication of cardiac ageing. Furthermore, we have shown that decorin induces microvascular inflammation, endothelial barrier dysfunction and diastolic dysfunction in the heart. Mechanistically, we propose that decorin acts as a ligand of the TLR2 receptor driving the expression of different pro-inflammatory cytokines like IL-1β in the mouse heart.

## Limitations

Although our study offers valuable insights into the role of decorin in the context of cardiac aging, our model does not proof a causal role of decorin in cardiac aging. In order to study what are the effects of decorin on the heart, we have used a systemic infusion approach using osmotic mini- pumps. This approach could induce systemic effects in the body, with secondary effects on the heart. Furthermore, the systemic application of recombinant decorin does not fully recapitulates the endothelial-specific induction, which we have observed in the aging heart. Finally, with our experiments, we cannot distinguish between potential intracellular effects and autocrine or paracrine activity of decorin. Nevertheless, our in vitro and loss-of-function studies support that decorin induces endothelial cell inflammation.

## Author contribution

G.L., Conceptualization, Data curation, Formal analysis, Investigation, Funding acquisition, Writing—original draft, Project administration. T.W., C.B., B.N.T, M.S., M.M-R., A.F., Investigation, Methodology. D.J., M.R-J, W.T.A. Single nucleus RNA sequencing. S.D. Conceptualization, Supervision, Writing—original draft, Project administration None of the authors have competing interests to declare

---

**Supplementary Figure 1.**
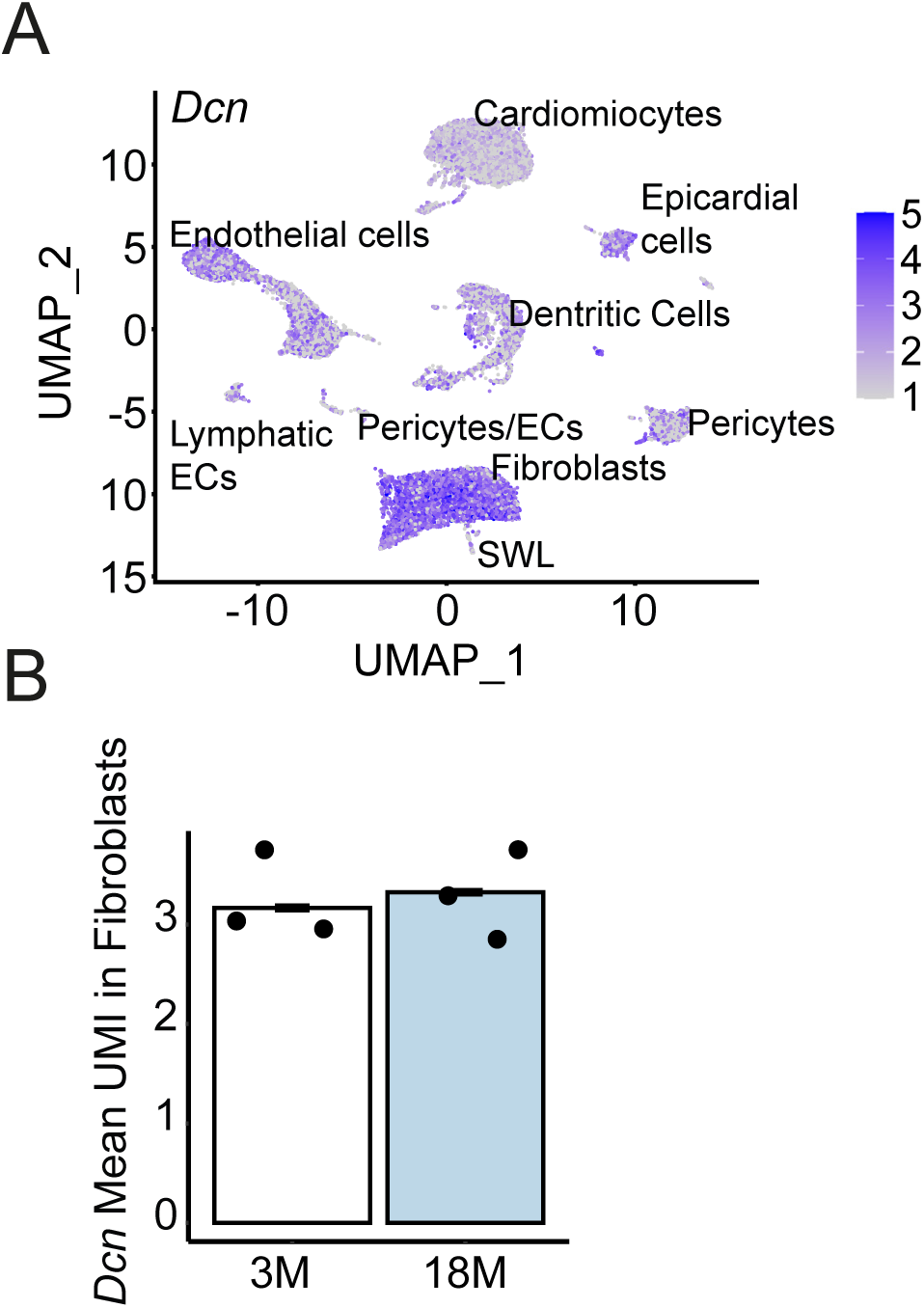
Decorin is expressed in fibroblasts but not regulated in ageing. Single nucleus RNA sequencing of young and old hearts. N=3. (**A**) UMAP plot depicting *Dcn* expression in the different cardiac cell types. (ECs, endothelial cells; SWL, Swan like cells) (**B**) Bar graphs depicting *Dcn* mean UMI expression in fibroblasts.

**Supplementary Figure 2.**
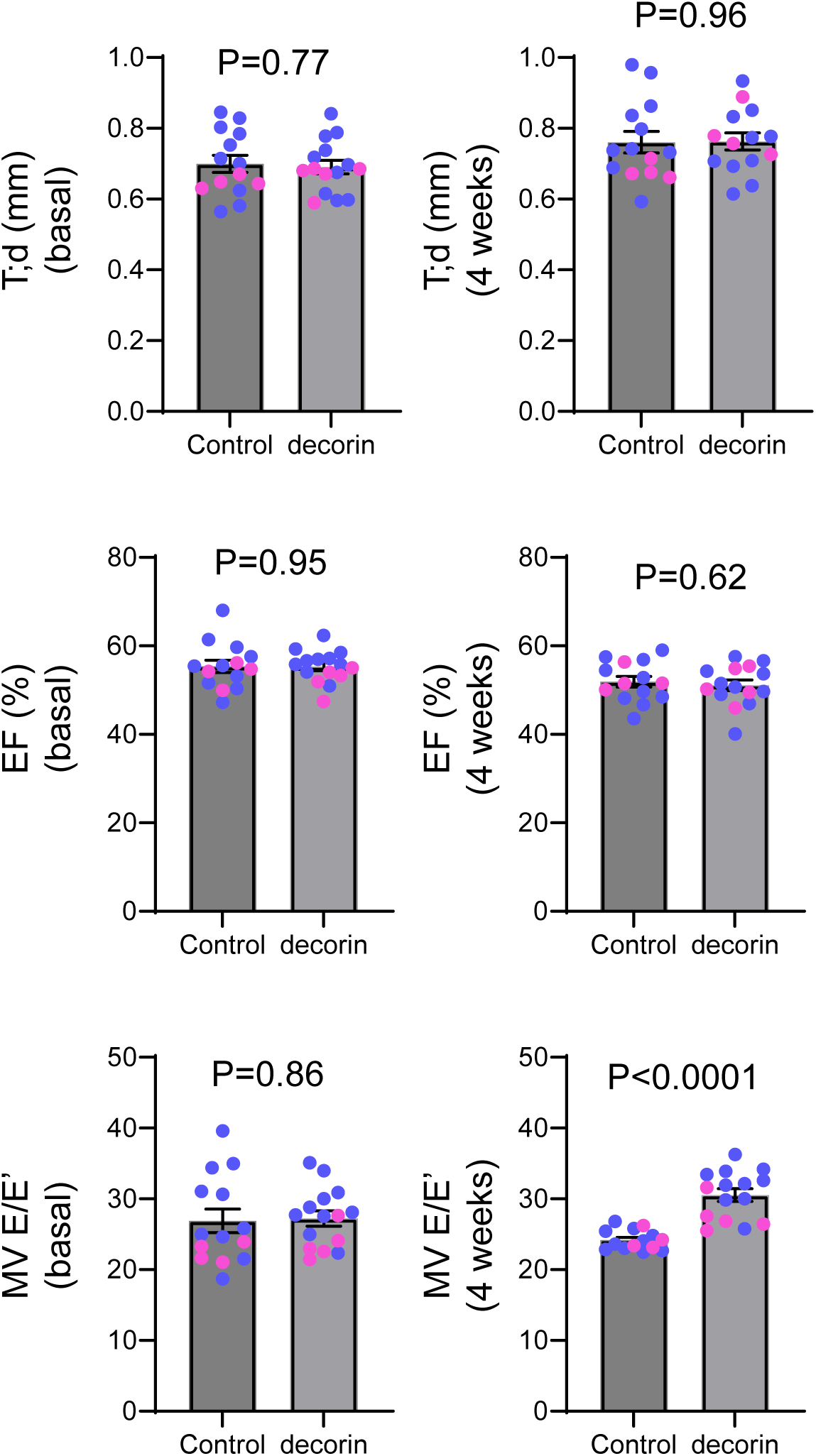
**Echocardiography absolute values.** Echocardiographic analysis of animals after decorin treatment. (N=14 for Control and N=15 for decorin). Colour indicates the sex of the animal, blue represents male, pink represents female. Data are shown as mean ± SEM. P-value was calculated by unpaired two-tailed Student’s t-test.

**Supplementary Figure 3.**
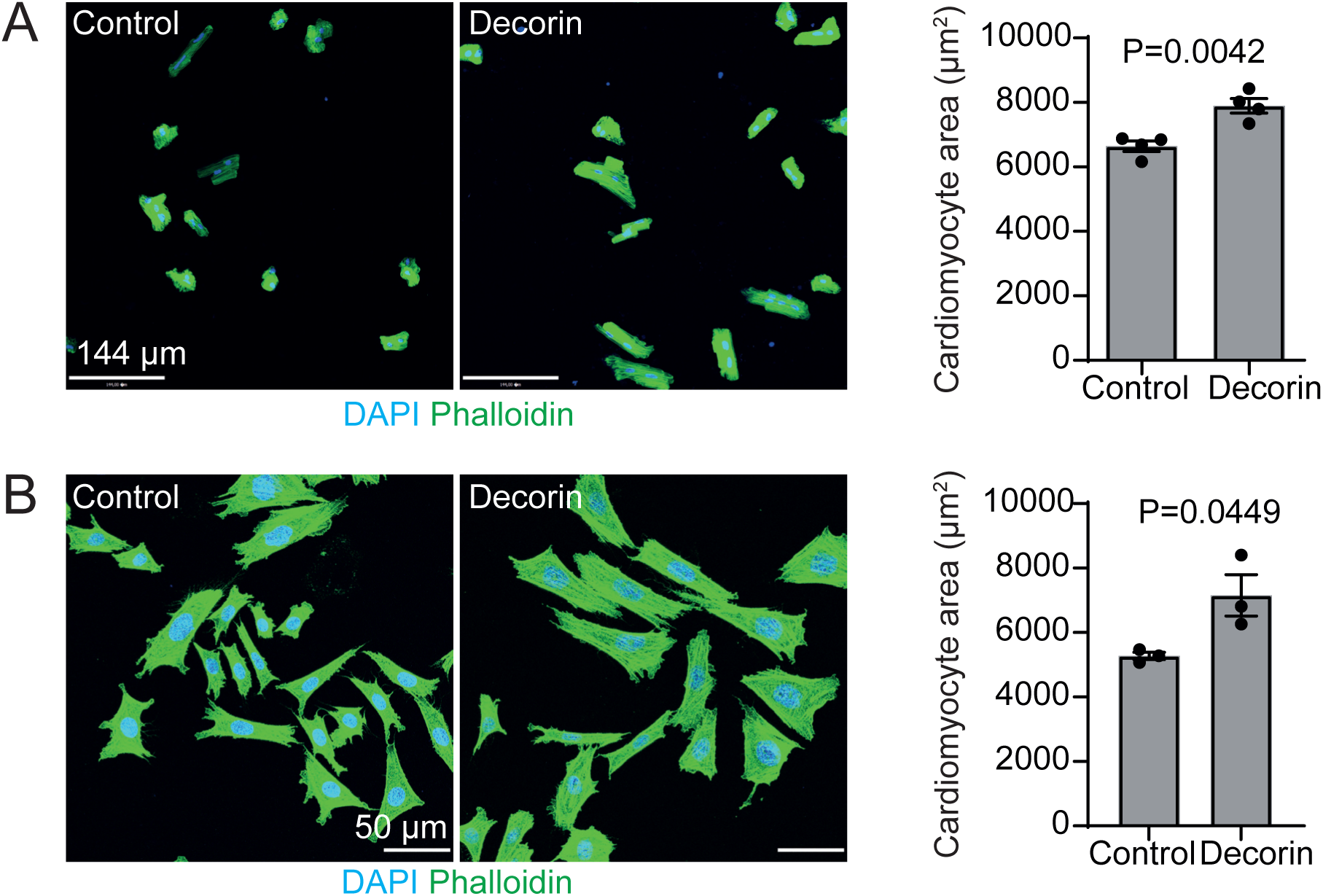
Decorin induced hypertrophy *in vitro*. (**A**) Immunostaining of freshly isolated murine cardiomyocytes. Treated cardiomyocytes are bigger than the control counterparts (N=4). Data are shown as mean ± SEM. P-value was calculated by unpaired two-tailed Student’s t-test. (**B**) Immunostaining of primary human cardiomyocytes. Treated cells are bigger than the control counterparts (N=3). Data are shown as mean ± SEM. P-value was calculated by unpaired two-tailed Student’s t-test.

**Supplementary Figure 4.**
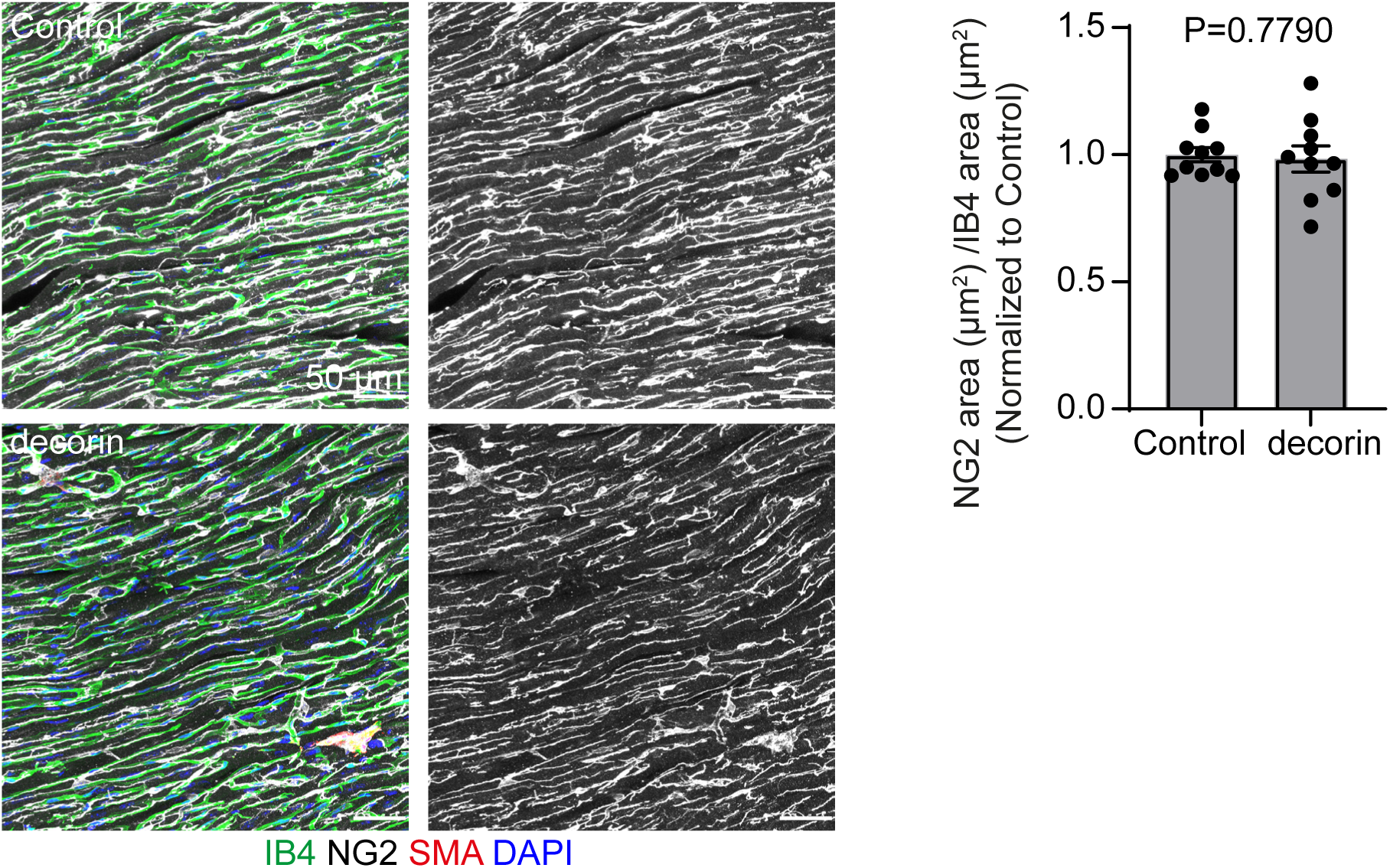
Pericyte coverage is not altered upon Decorin treatment. Immunostaining of cross sections through the left ventricle. Decorin treatment does not affect pericyte coverage (N=10). Data are shown as mean ± SEM. P-value was calculated by unpaired two-tailed Student’s t-test.

**Supplementary Figure 5.**
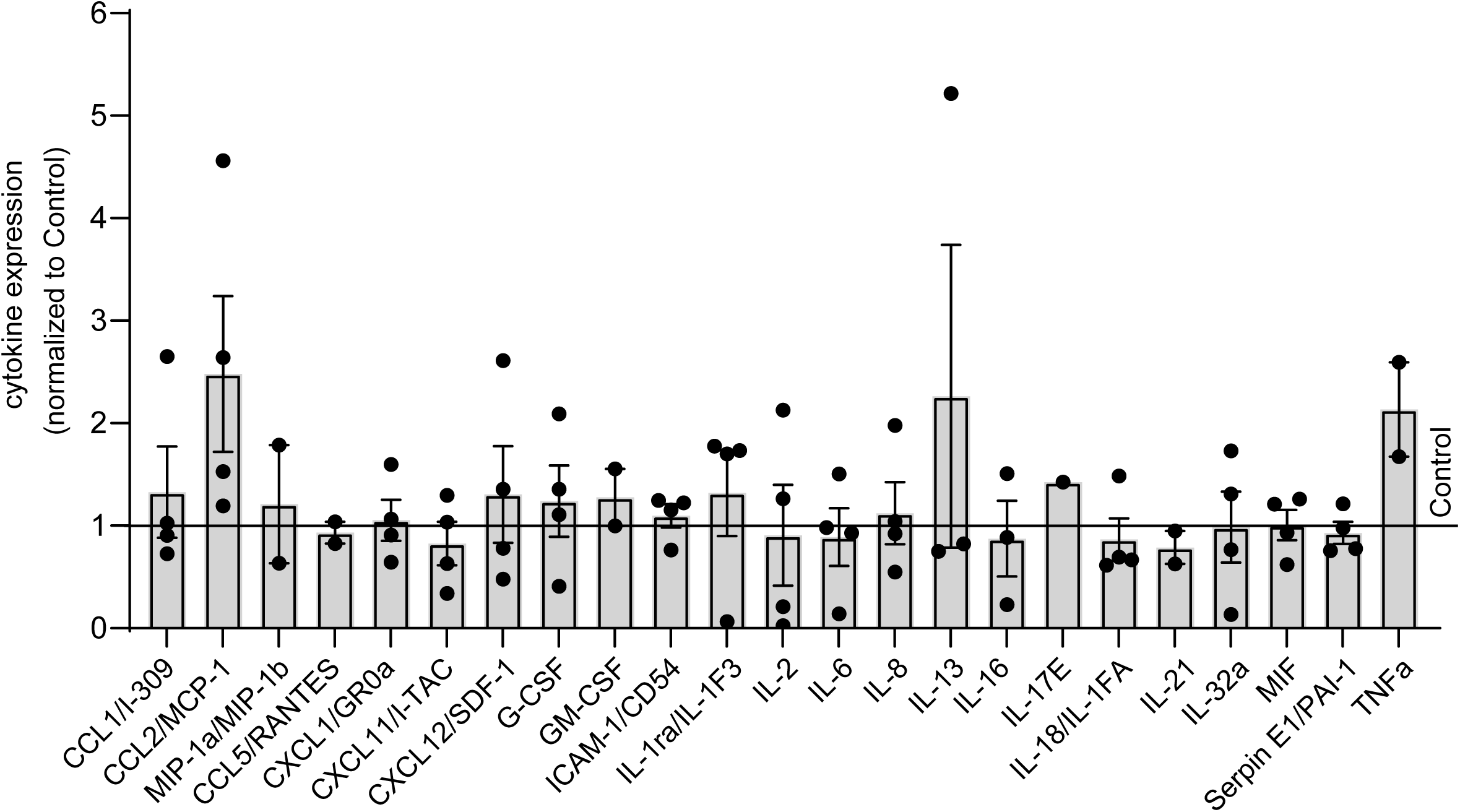
Complete cytokine array. Cytokine array expression analysis of HMVEC-Cs upon decorin treatment. Data represented as mean ± SEM. The horizontal line represents Control.

**Supplementary Figure 6.**
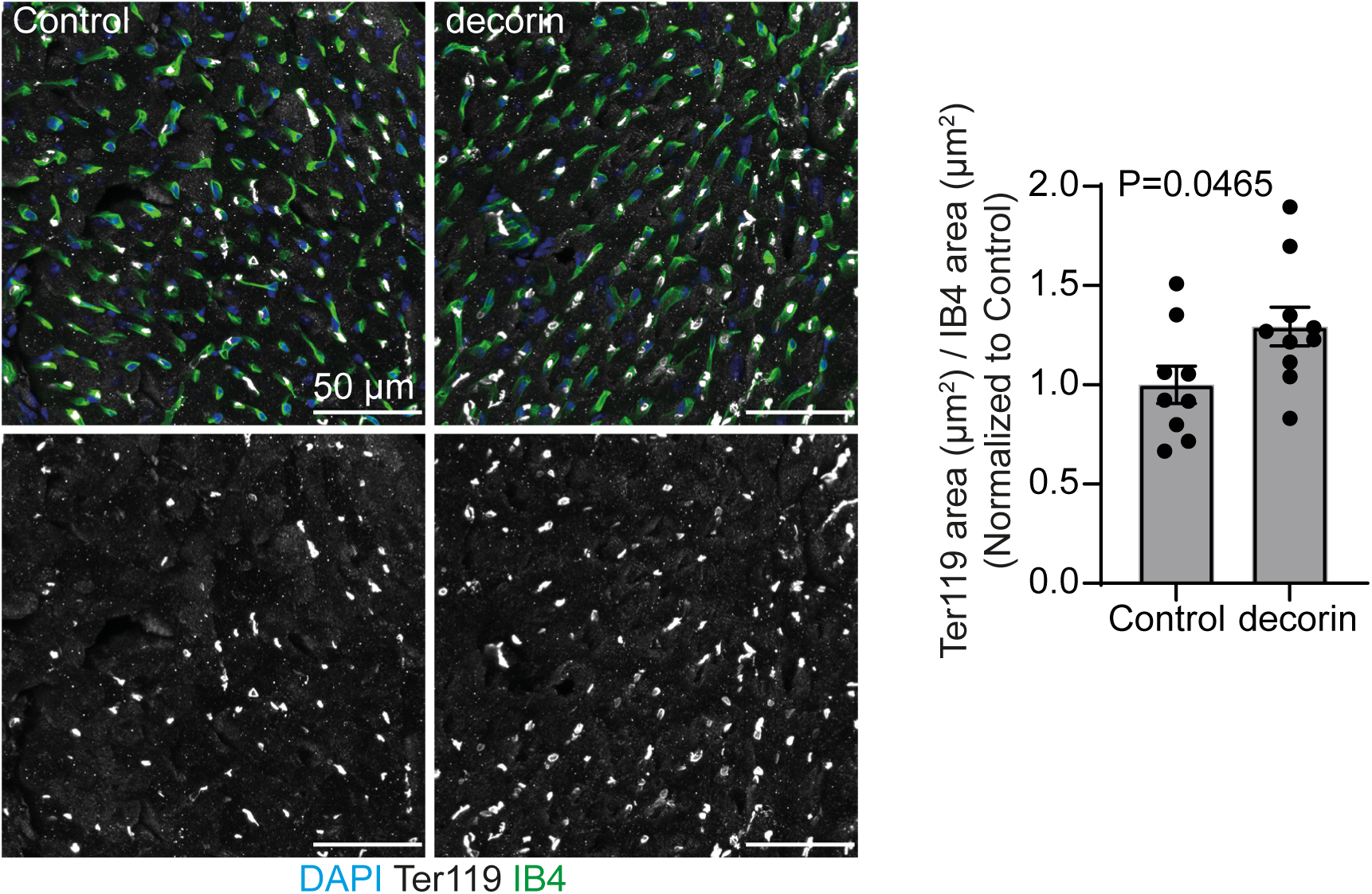
Decorin treatment induces erythrocyte accumulation. Immunostaining of heart cross sections (N=10). Decorin induces erythrocyte accumulation in the heart capillaries. Data are shown as mean ± SEM. P-value was calculated by unpaired two-tailed Student’s t-test.

